# Comparative network pharmacology of artificial sweeteners to understand Its health consequences

**DOI:** 10.1101/2024.03.29.587332

**Authors:** Gohit Tankala, Arun HS Kumar

## Abstract

**Background:** Artificial sweeteners (ASwt) are widely consumed sugar substitutes, but their long-term health effects remain a subject of debate. While regulatory bodies generally consider them safe at recommended doses, concerns persist regarding potential adverse effects. This study aimed to investigate the interactions between ASwt and biological targets using in silico analysis, focusing on target affinity, selectivity, and tissue expression.

**Methods:** Five common ASwt – acesulfame K (Ac), aspartame (As), sucralose (Su), steviol (St), and saccharin (Sa) were evaluated. Their target interactions were predicted using a cheminformatics approach, analysing affinity towards functional groups and protein targets. Concentration/affinity (C/A) ratios were calculated to assess the likelihood of target activation at achievable doses. Expression of high-affinity targets with significant C/A ratios in various organs was assessed using the Human Protein Atlas database. **Results**: The ASwt displayed potential to modulate most of the functional groups at physiologically feasible affinities. Ac exhibited a broad range of targets, while St showed a preference for kinases and proteases. Notably, As and Su demonstrated interactions with membrane receptors and kinases. C/A ratio analysis revealed potential concerns for As and Su. Several of its targets, including ROCK2, ACE, ITGA2/5, PIM2, KDM5C, PIM1, SLC1A2, SETD2, CAPN1, LTA4H, MKNK2, HDAC1 and CDK, showed high C/A ratios, suggesting possible functional modulation at achievable intake levels. Organ specific expression analysis identified the endocrine, respiratory, renal, reproductive, central nervous, digestive, and musculoskeletal systems as a region particularly susceptible due to the high expression of high affinity targets linked to cell growth, extracellular matrix, epigenetic regulations, and inflammation. Interestingly, 30 tissues expressed high-affinity targets for both As and Su, while 14 tissues exclusively expressed targets for As. **Conclusion**: This study highlights the potential for ASwt to interact with various biological targets, particularly As and Su. The high C/A ratios of some As targets and the tissue-specific expression patterns suggest potential safety concerns that require in vivo validation.

## Introduction

Artificial sweeteners (ASwt) have become ubiquitous in our diet, offering a sugar-free alternative for weight management, diabetes control, and/or simply satisfying calorie conscious sweet tooth^1^. However, their safety and potential health concerns remain a topic of ongoing research. The increased use of ASwt as alternatives to sugar can be attributed to the extensive marketing efforts by manufacturers, and the increased prevalence of metabolic syndrome, diabetes, obesity, as well as other metabolic disorders, wherein ASwt are perceived as safer substitutes^2^. ASwt are also often recommended for individuals who are diabetic, obese/overweight or those who are trying to manage their weight, as they seek healthier alternatives to regular sugar. However, there is little evidence defending this claim that ASwt consumption has a beneficial effect on these patients^3^. Regulatory bodies like the FDA and EFSA have deemed commonly used ASwt as safe for human consumption at recommended intake levels^4^. These levels are established through rigorous evaluations considering factors like metabolism, absorption, and potential toxicity. Despite safety evaluations, several studies suggest potential health risks associated with chronic consumption of ASwt. A WHO systematic review^3^ revealed that replacement of ASwt with sugar does not provide a means for weight management in the long-term, and several studies have discovered a positive correlation between long term ASwt consumption and risk of developing cardiovascular disease^5-8^, type 2 diabetes mellitus^6,8-11^, and mortality in adults^6,8,9,12^. Current literature also reveals other concerning associations between ASwt consumption and various routinely observed clinical conditions, including heightened risks of developing cancer^13-15^, chronic kidney disease^16,17^, adiposity related diseases^8,9,18,19^, as well as non-alcoholic fatty liver disease^20^.

The adverse effects following chronic intake of ASwt can be consequence to disruptions to insulin signalling and gut microbiota, potentially influencing blood sugar control, impacting digestion, nutrient absorption, overall gut health and increasing the risk of metabolic syndrome^21,22^. Despite these reported clinical associations, there is substantial research gap regarding the pharmacodynamic effects of these sweeteners in homo sapiens, leading to lack of insights into their mechanisms of actions. Although ASwt offer the so-called “sugar-free option”, their long-term health effects require exploration of the pharmacological mechanisms of actions. Hence, research into the pharmacodynamics of ASwt can provide a foundation for establishing a mechanistic basis for highlighting safe consumption practices and mitigating potential health risks associated with their unaccounted consumption, perceiving them to the safe. A literature search revealed that the following were the top 5 most consumed ASwt in sugar-free food and beverages, Acesulfame K (Ac), Aspartame (As), Saccharin (Sa), Steviol (St), and Sucralose (Su), all of which are approved by the US-FDA, EFSA, and various other global food safety organisations for use as sweetening agents.^4,23-27^ The current literature gap on the pharmacodynamics properties of these ASwt, led us to plan this study to address the gap using a network pharmacology approach. Which will help us understand the receptor binding profiles of the different ASwt and therefore establish a foundational understanding of how they interact with the human body, potentially uncovering mechanisms behind currently observed health associations. This perspective offers insights into the molecular mechanisms that are underlying currently observed adverse health effects, as well as a possibility to highlight potential health risks associated with the consumption of ASwt.

## Materials and Methods

The isomeric SMILES sequence of each sweetener (Ac = “CC1=CC(=O) [N-]S(=O)(=O)O1”, As = “COC(=O)[C@H](CC1=CC=CC=C1)NC(=O) [C@H](CC(=O)O)N”, Sa = “C1=CC=C2C(=C1)C(=O)NS2(=O)=O”, St =“C[C@@]12CCC[C@@]([C@H]1CC[C@]34[C@H]2CC[C@](C3)(C(=C)C4)O)(C)C(=O)O”, Su = “C([C@@H]1[C@@H]([C@@H]([C@H]([C@H](O1)O[C@]2 ([C@H]([C@@H]([C@H](O2)CCl)O)O)CCl)O)O)Cl)O”) was acquired from the PubChem database, which was then inputted into Swiss Target Prediction software (http://www.swisstargetprediction.ch/) to predict and identify the targets of each sweetener, specific to homo sapiens. The 2D structure of the ASwt in the output files of Swiss Target Prediction software were used in this study to compare their structures. The commonalty of the targets and target classes between the ASwt were assessed using Venn Diagrams.^28^ The Uniprot database (https://www.uniprot.org/) was used to obtain the protein sequence of each individual target of the ASwt, and Yuel tool (https://dokhlab.med.psu.edu/cpi/#/YueL) and Autodock Vina 1.2.0 were used to predict the affinity between the sweeteners and each of their potential targets as described before.^29-33^ The targets were then classified into various functional groups, to assess the selectivity of each ASwt to any specific functional group(s).

The pharmacokinetic properties of the sweeteners were assessed using data reported in current literature. The volume of distribution (Vd) obtained from current literature^4^, and the dosage values (DV) which were obtained by evaluating current average daily intake (ADI) recommendations^4^ by the US-FDA, and EFSA, as well as current data regarding their consumption^23-27,34^ were trichotomized into low, medium, high ranges. The Vd and DV were used to calculate the effective plasma concentration (µM) achieved at the three different DV for each individual sweetener.

The Concentration/Affinity (C/A) ratio was calculated for each of the ASwt target, as a ratio of plasma concentration of the ASwt and its affinity value to its specific target. The C/A values obtained were used to generate a heatmap of each of the ASwt and their targets using the conditional formatting tool in Microsoft Excel software. C/A ratio ≥ 1.9 was used as a threshold to investigate the targets that are most likely to have an acute pharmacodynamic impact. This threshold was considered based on C/A ratio ≥ 1.9 accounting to an ASwt plasma concentration ∼ twice the value of its affinity to its target and hence most likely to have a pharmacodynamic effect.

Following identification of high affinity targets, the tissue specific expression of high affinity targets (based on C/A ratio ≥ 1.9) was individually assessed using the ProteinAtlas database (https://www.proteinatlas.org/), and classified protein expression into the following three categories; “Not detected, Low, Medium, High”. If the protein expression was unavailable, the RNA expression was assessed and the following ranges were used; High = 70-100% nTPM, Medium = 40-69% nTPM, Low = ≤ 39% nTPM. If the target’s tissue specific expression was classified as either “Not Detected” or “Low”, they were excluded from our further analysis.

## Results

The 2D structures of all the five ASwt are presented in figure 1. The defined dosage values (DV; mg/day) were calculated to be as follows in the order of low, medium, and high; [Ac (450, 900, 2000), As (1000, 2500, 5000), Sa (150, 300, 600), St (100, 240, 500), Su (150, 300, 600)] (Figure 1). The following Vd (L) values of each sweetener was estimated using data reported in literature; Ac (110), As (109), Sa (264), St (100), Su (100), and was used to predict the effective plasma concentration (mg/L) achievable in humans at each DV and were calculated to be as follows in the order of low, medium and high; [Ac (4.09, 8.18, 18.18), As (9.17, 22.94, 45.87), Sa (0.57, 1.14, 2.27), St (1, 2.4, 5), Su (1.5, 3, 6)] (Figure 1).

**Figure 1:**
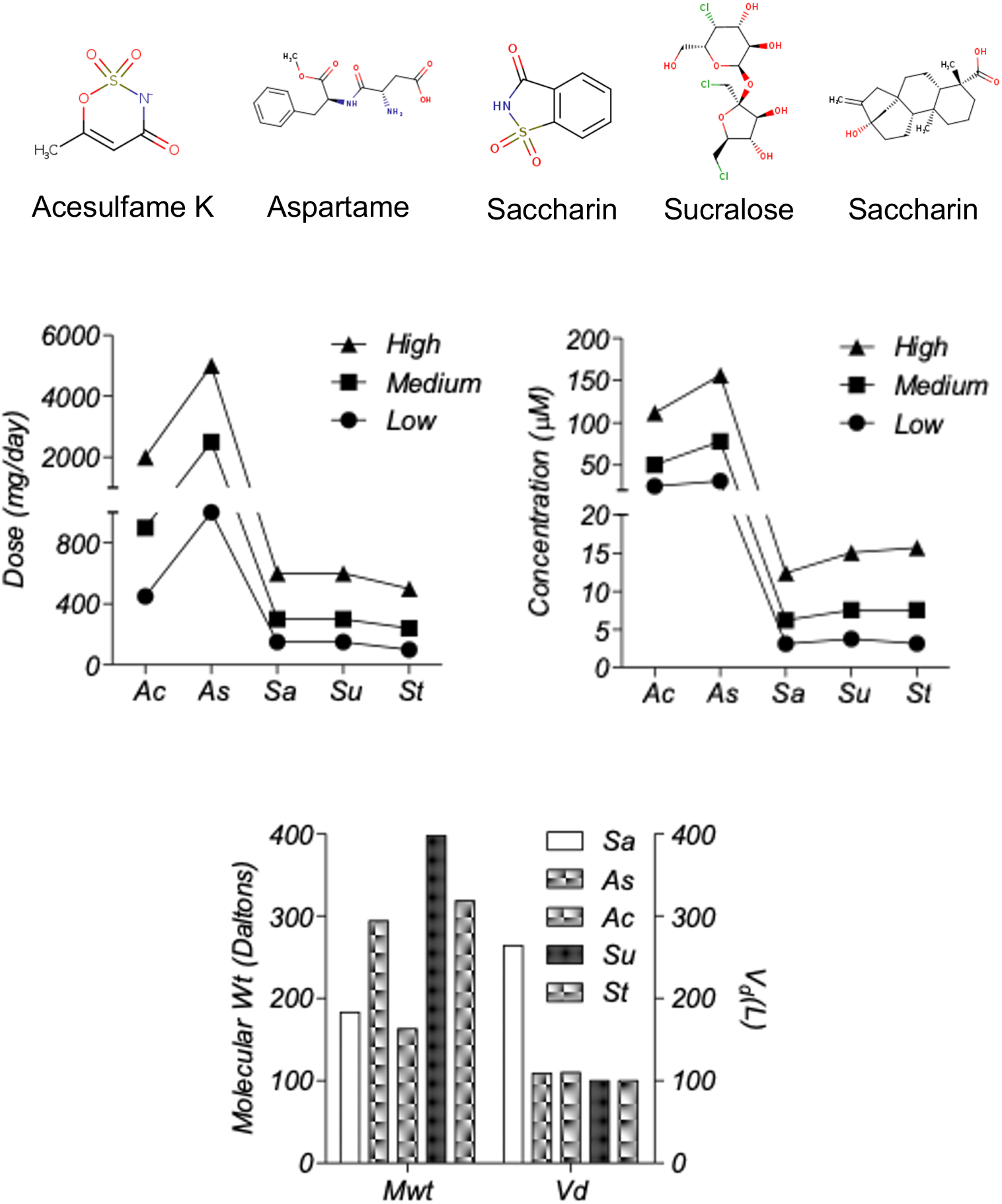
Pharmacological Properties of the different artificial sweeteners (ASwt). Chemical Structures of the different Artificial Sweeteners (ASwt). The middle graphs show the trichotomized (low, medium, high) data of ASwt doses (mg/day) and predicted plasma concentration (µM), in humans. The bottom graph shows the molecular weight (Daltons), and the volume of distribution (L), of each ASwt.

These values were then converted into µM by using the molecular weight (Daltons) of each sweetener; Ac (163.15), As (294.31), Sa (183.19), St (318.4), Su (397.63), and were calculated to be as follows in the order low, medium and high; [Ac (25.1, 50.2, 111.54), As (31.21, 78.01, 156.03), Sa (3.1, 6.21, 12.42), St (3.14, 7.55, 15.72), Su (3.77, 7.54, 15.08)] (Figure 1).

Collectively the ASwt were observed to target 23 different functional groups. To evaluate if the sweeteners had any selectivity to specific functional groups, the mean affinity of each sweetener to their respective functional groups were assessed (Figure 2). Most of the functional groups were targeted by ASwt at physiologically feasible affinities (<1000 µM). The highest affinity of Ac was discovered to be towards erasers and enzymes, whilst the least affinity was towards proteases and lyases. As had the greatest affinity towards electrochemical transporters, oxidoreductases, writers, and erasers whilst the least affinity was discovered to be towards surface antigens, proteases, and enzymes. Sa showed selectivity towards transporters, kinases, cytosolic proteins, and ion channels whilst they showed the least affinity towards lyases, nuclear receptors, and family A G-Protein Coupled Receptors (GPCR). St had the greatest affinity towards kinases, proteases, secreted proteins, oxidoreductases, and membrane receptors, whilst it was revealed that they showed the least affinity towards fatty acid binding proteins, cytosolic proteins, and phosphatases.

**Figure 2:**
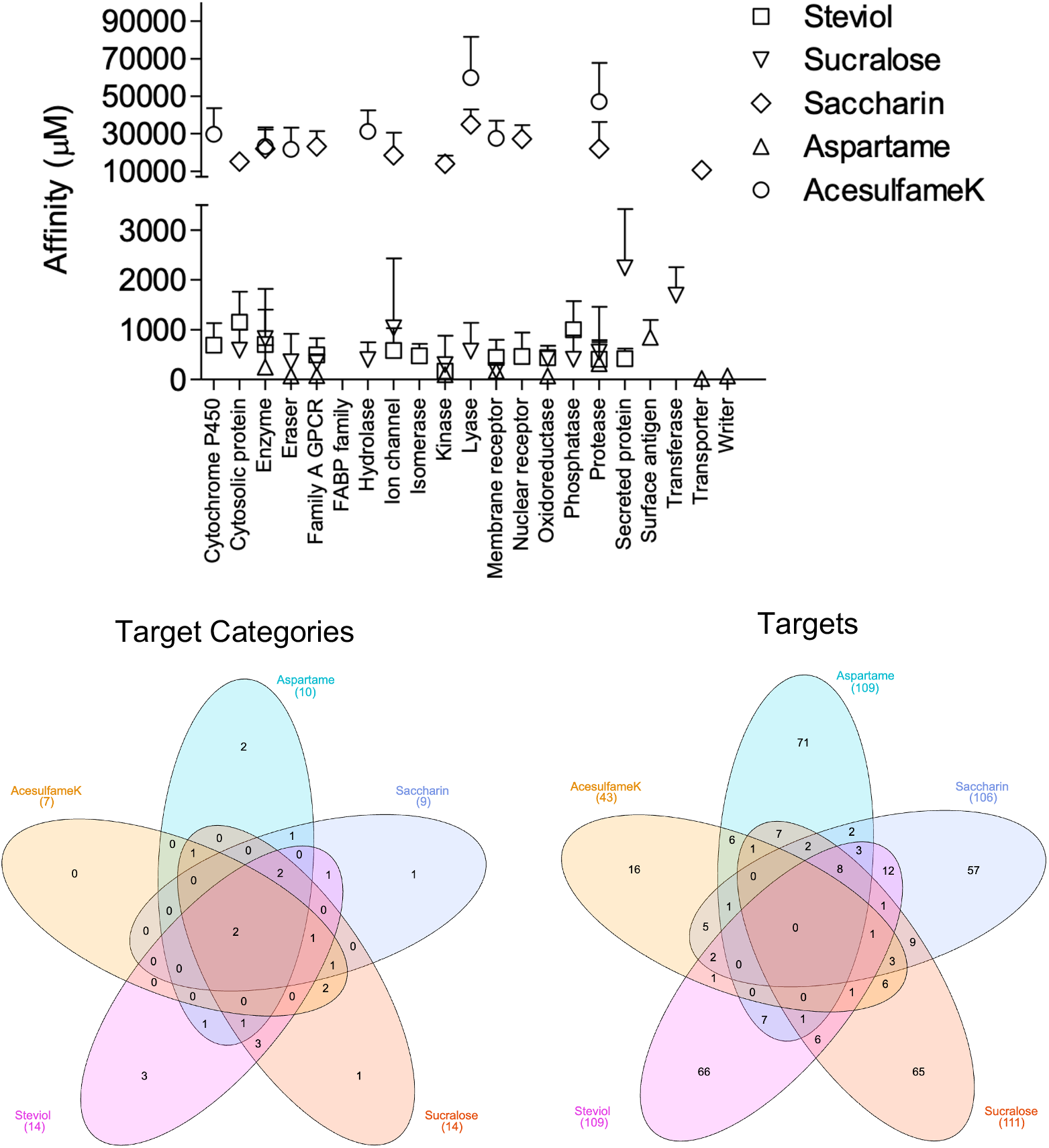
Affinity of artificial sweeteners (ASwt) to functional groups. The graph shows the affinity (µM, mean ± SD) of all 5 ASwt to various functional groups; Steviol =□, Sucralose = △, Saccharin = ⍰, Aspartame = ⍰, Acesulfame K = ⍰. The 1^st^ Venn Diagram is comparison of the different ASwt targeting their different functional groups, and the 2^nd^ Venn Diagram is comparing the different ASwt and their predicted targets.

The highest affinity of Su was revealed to be towards membrane receptors, kinases, family A GPCR, erasers, hydrolases, and phosphatases, whilst they showed the least affinity towards secreted proteins, transferases, and ion channels.

Venn Diagrams^28^ were used to examine if different ASwt shared common functional groups as their targets (Figure 2). Enzymes, and proteases were common targets of all 5 ASwt. Ac, Sa, St and Su had cytosolic proteins as their common targets. As, Sa, St and Su shared Family A GPCR, and kinases as common targets. Membrane receptors were common targets among Ac, As, St and Su. Oxidoreductases were common targets of As, St, Su, while erasers were common targets of Ac, As and Su. Lyases were common targets of Ac, Sa and Su. Ion channels were common targets of Sa, St and Su. Nuclear receptors were common targets of Sa and St. Phosphatases, and secreted proteins were common targets of St and Su. Hydrolases were common targets of Ac and Su. The exclusive functional groups of As were electrochemical transporters, surface antigens, and writers. Sa had one exclusive functional group i.e., transporters. St had following three exclusive functional groups i.e., cytochrome P450, fatty acid binding protein family, and isomerases. The exclusive functional group of Su was transferase. Except for Ac, all other ASwt exclusively targeted 1-3 different functional groups (Figure 2).

The network analysis identified potential targets of all the different ASwt i.e., Ac (43), As (109), Sa (106), St (109), Su (111). Venn Diagrams^28^ were again used to examine if different ASwt shared their targets (Figure 2). Notably, 8 targets were shared by As, Sa, St and Su, i.e., PSENEN, PTGS2, ACE, PSEN1, PSEN2, APH1B, NCTSN, APH1A. Ac, Sa, St and Su shared 1 target; MCL1, whilst Sa, St and Su shared PTGES. Ac, Sa and Su shared CA9, CA2, and ELANE. Ac, St and Su shared GSK3B. Ac, Sa and St shared CES2, and BCHE. Ac, As and Sa shared MMP2, whilst Ac, As and Su shared CASP3. As, Sa and St shared NOS2, OPRD1 and OPRM1, whilst As, Sa and Su shared PIM1 and F2. As, Su and St shared EDNRA, and Sa, St and Su shared PTGES. Ac and Su shared IDO1, TYMP, CDC25B, TYMS, CASP6, and CASP7, whilst Ac and St shared SIGMAR1. Ac and Sa shared STAT3, CES1, AOC3, CA12, and CA1 whilst Ac and As shared NAAA, KDM5C, KDM5B, KDM4A, KDM4C and KDM4B. As and Sa shared HDAC1 and OPRK1. As and Su shared KDM5A, CDK2, CSNK2A1, ITGAL, ITGB2, ICAM1, and DPP4. As and St shared AURKA, ITGA4, ITGB1, HMGCR, MME, BACE1, CPA1. Sa and St shared HSD11B2, HSD11B1, HTR2A, PTGER2, DRD5, AGTR1, PTGDR2, BCL2, PPARG, NR1H3, RORC, and F10. Sa and Su shared HSP90AA1, GBA, AKR1C3, FBP1, AKR1B1, P4HTM, MMP1, MMP3 and MMP9, whilst Su and St shared ADORA3, P2RX3, CDC25A, PTPN2, and PTPN1. The exclusive targets of Ac were CYP2A6, PADI2, PADI1, PADI3, PARP1, PADI4, DPYD, XDH, KAT2B, SIRT2, SIRT3, GDA, ACHE, TLR9, CASP9 and CASP4. The exclusive targets of As were SLC1A2, SLC5A1, SLC1A3, SLC15A1, SLC6A3, PAM FNTA, TDP1, NOS3, BIRC3, BIRC2, YARS, FNTB, CBX4, PPIA, HDAC8, NTSR1, TACR2, GHSR, TACR1, AGTR2, OXTR, TACR3, S1PR5, GALR1, FPR1, CALCRL, ROCK2,, PIM2, MKNK2, ILK, EPHA2, MAPKAPK2, ITGA5, ITGA2, ITGA2B, ITGAV, IL2RA, ITGB3, RXRA, XIAP, PDYN, IL1B, TUBB1, RRM1, CAPN1, LTA4H, RNPEP, DPP8, REN, BACE2, CASP1, PGC, CTSE, ANPEP, CTSD, LAP3, CELA1, CPB1, TPSAB1, KLK5, CTRB1, CTRC, CBX7, HLA-A, HLA-DRB3, HLA-DRB1, SETD2, PRMT1, CARM1 and SETD7. The exclusive targets of Sa were discovered to be SOAT1, DAGLA, PTPRC, NAMPT, HSD17B2, HSD17B1, AKR1C1, SERPINE1, ADRA1A, UTS2R, HTR2C, ADRA1D, CCR8, ADRA1B, CNR1, LPAR2, GPR55, CHRM1, CNR2,, CHRM3, HTR1B, HRH3, HTR1A, DRD3, DRD4, DRD2, GRIN2B, GRIN1, BCL2L1, SCN9A, TRPV4, KCNA5, TRPA1, KCNH2, ABL1, LIMK1, ERBB2, IKBKB, LIMK2, CA5A, CA13, CA5B, CA4, CA6, CA14, CA7, NR3C2, NR1H2, BMP1, MMP8, ADAMTS4, MMP13, EPHX1, PLAU, PRSS1, SLC6A4 and SLC22A6. The exclusive targets of St were revealed to be CYP17A1, CYP51A1, CYP26A1, CYP26B1, KIF11, BCL2L2, TP53, BCL2A1, SCD, UGT2B7, G6PD, AMPD2, TERT, PLA2G4A, HSD17B3, AMPD3, CPT2, FAAH, AMPD1, UBA2, POLB, SAE1, AKR1B10, PLA2G1B, CPT1B, HCAR2, GPBAR1, PTGFR, EDNRB, CCKBR, GABBR1, FABP2, FABP4, FABP1, GABRB2, GABRG2, GABRA2, TOP2A, TOP1, PRKCH, FLT1, GSK3A, FGFR1, NPC1L1, CD81, MDM2, THRA, AR, ESR2, THRB, PPARD, RARA, NR1H4, PPARA, RARG, VDR, IMPDH2, CDC25C, PTPN6, ACP1, PREP, CTSA, EPHX2, ECE1, SERPINA6 and SHBG.

The exclusive targets of Su were found to be HSPA8, LGALS9, LGALS8, LGALS4, HSPA5, PDCD4, HEXA, AHCY, PYGL, TREH, HEXB,, HK2, PYGB, PYGM, OGA, FUCA1, HK1, AMD1, ADK, HPRT1, DAO, PNP, HPSE, AKR1C2, CDA, PIN1, JMJD1C, KDM4E, ADORA1, ADORA2A, ADORA2B, GAA, AMY2A, ADA, TRPV1, CDK9, CCNA2, CCNT1, CCNB1, CDK4, CDK1, EGFR, GRK1, LCK, FYN, MAPK1, CCND1, CCNA1, SLC5A2, GAPDH, TYR, KMO, PTPRB, PTPN11, FOLH1, CASP2, NAALAD2, ADAM17, CASP8, GGH, FGF1, VEGFA, FGF2, DTYMK and TK1.

The affinity of Ac to its targets ranged from 4364.28 µM to 83111.14 µM, of which the high affinity targets were PADI2 (4364 µM), KAT2B (8305.64 µM), and KDM5C (9032.40 µM) (Figure 3). However, none of these potential targets had a significant C/A ratio (≤ 0.026). Our analysis revealed As to have 58 potential targets with a significant C/A ratio (≥ 1.9), of which the following 8 targets had an alarming C/A ratio ≥ 20; ROCK2, ACE, ITGA5, PIM2, KDM5C, PIM1, SLC1A2, SETD2. The highest affinity recorded was for ROCK2 (0.1806 µM) which had a C/A ratio of 838.577 for the high dose value (Figure 3). The affinity of Sa to its targets ranged from 3361.6 µM to 69202.9 µM, of which the highest C/A ratio was determined to be 0.004 and all its targets were therefore deemed insignificant as it is unlikely to achieve concentrations sufficient to activate these targets (Figure 3). The affinity of St to its targets ranged from 52.5 µM to 8910.5 µM, of which the high affinity targets were GSK3B (52.5 µM), ACE (52.7 µM), PRKCH (54.6 µM) (Figure 3). However, none of these potential targets had a significant C/A ratio (≤ 0.299). Su was found to have 5 potential targets with a significant C/A ratio (≥ 1.9); CDK4, CDK9, SLC5A2, CDK1, EGFR (Figure 3). CDK4 had the highest affinity (0.60 µM) and a C/A ratio of 24.999 at the high dose value (Figure 3).

**Figure 3:**
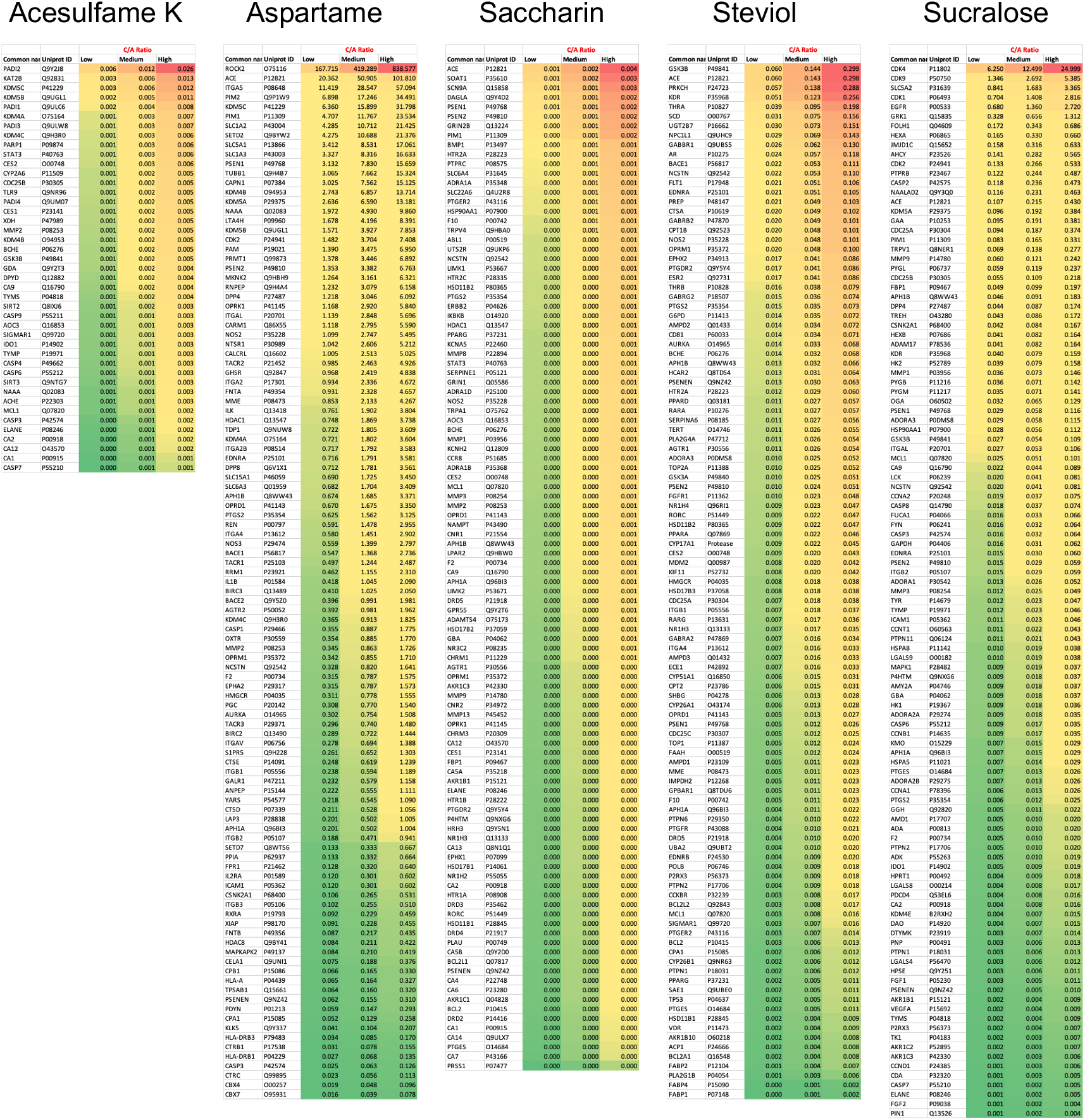
Concentration/Affinity (C/A) Ratio of the 5 artificial sweeteners (ASwt) to their respective predicted targets. The heat maps represent the C/A Ratio of the ASwt against all of its identified targets from the SwissTargetPrediction database, at each trichotomized dosage value (low, medium, high) (Scale: Red to Green = High to Low C/A ratio values).

To assess the organ specific impact of ASwt, we examined an organ specific expression pattern of the high affinity targets with a focus on the significant targets of As (58) and Su (5), whilst the targets of Ac, Sa, St were excluded from this part of the study based on the low C/A ratio (Figure 4). In the human Protein Atlas database, we identified 56 different organ types expressing targets of As and Su. The expression of the targets was classified as either high (green), medium (red), or low/none (blank) (Figure 4). We further defined these targets as highly significant if the target was highly expressed in > 15 organs, and the resultant highly significant targets identified were as follows: CAPN1 (30), LTA4H (16), MKNK2 (25), ITGA2 (17), HDAC1 (19), CDK9 (25). Of these targets CAPN1, LTA4H, MKNK2, ITGA2, HDAC1 are targets of As, and CDK9 is the only one that’s a target of Su. To define which organs were most likely to be pharmacodynamically affected, we focused on the organs that highly expressed the high affinity targets we had initially defined as significant (C/A ≥ 1.9). Forty-four organs were identified to express high affinity targets of As and Su (Figure 4). If a tissue highly expressed the target ≥ 10 times, we defined it as pharmacodynamically significant, and the organs we discovered to fit these criteria were colon, duodenum, kidney, placenta, rectum, small intestine, stomach, testis, cerebral cortex, cerebellum, bone marrow, appendix and tonsil (Figure 4). The expression of various high affinity targets of As and Su in various organs is also summarised in the bottom panel of figure 4. While 30 tissues had high expression for high affinity targets of both As and Su, and 14 tissues exclusively expressed high affinity targets of As (Venn diagram Figure 4). The organ systems which can be preferentially targeted by ASwt were endocrine, respiratory, renal, reproductive, central nervous, digestive, and musculoskeletal systems.

**Figure 4:**
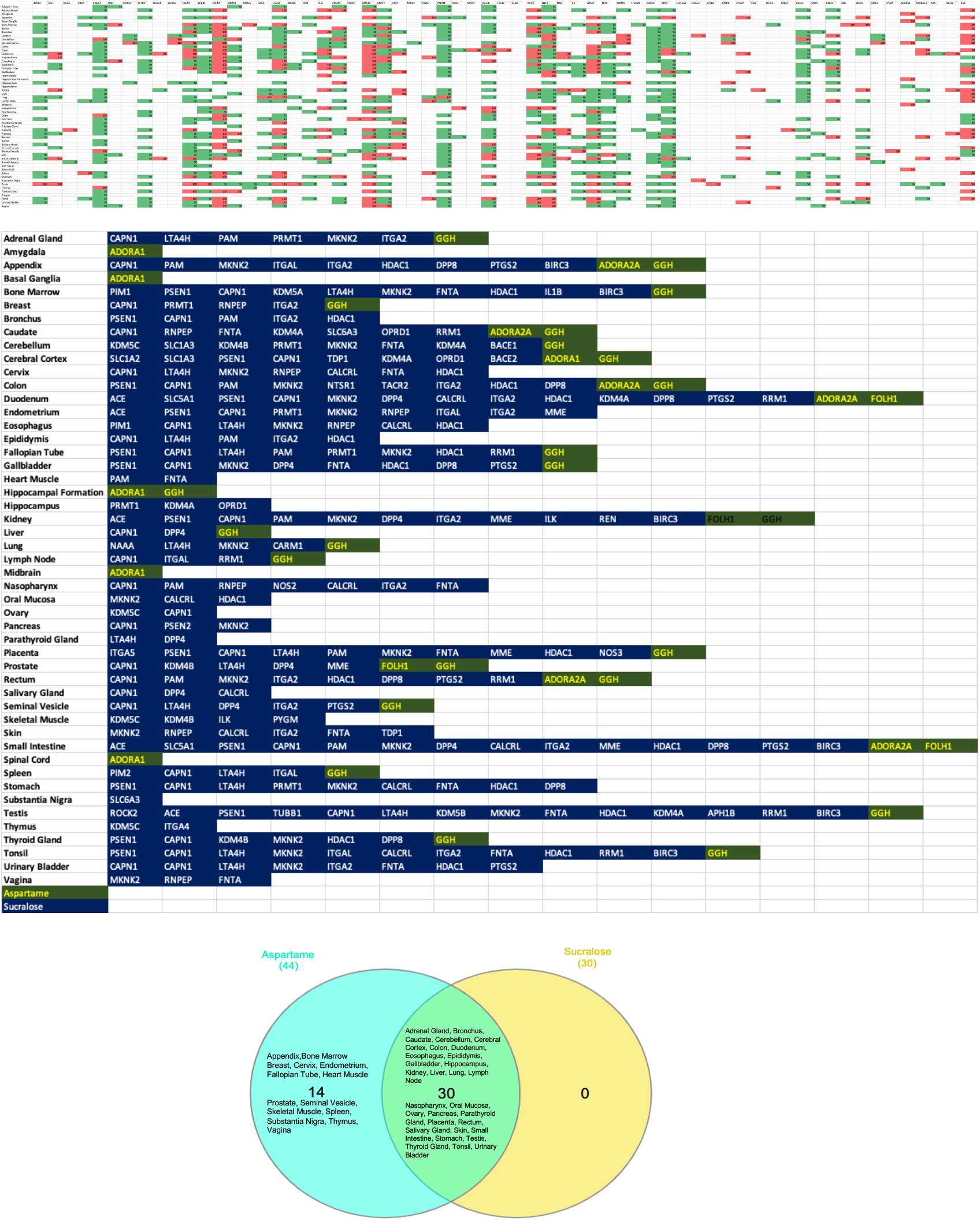
Organ specific expression analysis of significant artificial sweeteners’ (ASwt) targets. The upper graph shows the significant ASwt targets being graphed against different tissues. (Red = High Expression, Green = Medium Expression, Blank = Low/No Expression). The bottom graph summarises the organs highly expressing the significant targets of aspartame (blue) and sucralose (green), showing the different organs and the high affinity targets. (Blue = Aspartame Targets, Green = Sucralose Targets). The Venn diagram compares the organs expressing high affinity targets between aspartame (light blue) and sucralose (Yellow).

## Discussion

This in silico study is the first of its kind which investigated the potential interactions between five common artificial sweeteners (ASwt) and various biological targets. Our findings shed light on potential mechanisms by which ASwt may exert pharmacodynamic effects in humans. The network pharmacology approach has revealed several potential mechanistic insights that may explain currently observed associations between ASwt consumption and the development of various clinical conditions including cardiovascular disease, lipid disorders, endocrine disorders, type 2 diabetes mellitus, chronic kidney disease, and cancer. This wide range of disease risks associated with ASwt consumption are consistent with the diverse organ systems (endocrine, respiratory, renal, reproductive, central nervous, digestive, and musculoskeletal systems) targeted with high affinity by ASwt. Our study also highlights the dissimilarity between different ASwt examined in this study regarding their safety and pharmacodynamic effects, which in our view should influence safe consumption practises. Specifically, As and Su were identified to be least safe ASwt, based on their target profile and associated C/A ratios. While Sa was identified to be most safe ASwt followed by Ac and St based on their target profile and associated C/A ratios.

The famous quote by Paracelsus “only the dose makes a thing a poison”^35^ becomes very relevant to specifying safe consumption levels of ASwt. This principle is the foundation of safety, recognizing that any substance, even water or oxygen, can be harmful at high enough concentrations. While regulatory bodies have established safe intake levels for each ASwt^4^, our findings suggest potential reasons for re-evaluation, particularly for As and Su. Our analysis revealed that As and Su will interact with cellular targets at achievable doses, raising concerns about potential health consequences. This is especially relevant considering the high C/A ratios observed for some of its targets. Therefore, it is crucial to emphasize the importance of adhering to recommended intake levels and to consider the potential cumulative effects when consuming ASwt products. Long-term studies are necessary to definitively determine the safety of chronic ASwt consumption at these recommended doses. Nevertheless, based on our observations in this study, we suggest limiting daily intake levels of As and Su under 400 mg/day and 100 mg/day respectively or alternatively considering using Sa, Ac and St with daily intake limited to 150 mg/day, 400 mg/day and 80 mg/day respectively. These revised daily intake suggestions should be considered while designing long-term randomised clinical trials.

A considerable structural difference is also evident between different ASwt. Ac is a sulfamate ester that is 1,2,3-oxathiazin-4(3H)-one 2,2-dioxide substituted by a methyl group at position 6.^36^ As is the methyl ester of the aspartic acid and phenylalanine dipeptide.^36^ Sa is a 1,2 benzisothiazole with a keto group at position 3 and two oxo substituents at position 1.^36^ St is the basic backbone of steviol glycosides such as stevioside, rebaudioside A which are extracted from the stevia plant, it is a diterpene compound that consists of a tetracyclic diterpene structure featuring a lactone ring, with a hydroxyl group located at position 13.^36^ Su is a disaccharide derivative composed of 1,6-dichloro-1,6 dideoxyfructose and 4-chloro-4-deoxygalactose, produced by the chlorination of sucrose which results in three chlorine atoms replacing three hydroxyl groups, thereby preventing it from being broken down.^36^ These structural differences may account for the considerable variations in their targets/ target groups observed in this study. The comparative analysis of the various functional groups of the targets impacted by ASwt allows us to assess how the most used combinations of ASwt can influence systemic physiology. The most used combinations of ASwt in artificially sweetened beverages are as follows; Ac and As (Coke Zero, 7up Zero), As and Sa (Fountain Diet Coke)^37^, Ac and Su (Red Bull Sugar Free). The Ac and As combination binds with 13/28 functional groups (Cytosolic Protein, Electrochemical Transporter, Enzyme, Eraser, Family A GPCR, Hydrolase, Kinase, Lyase, Membrane Receptor, Oxidoreductase, Protease, Surface Antigen and Writer). The As and Sa combination binds with 15/28 functional groups (Cytosolic Protein, Electrochemical Transporter, Enzyme, Eraser, Family A GPCR, Ion Channel, Kinase, Lyase, Membrane Receptor, Nuclear Receptor, Oxidoreductase, Protease, Surface Antigen, Transporter, and Writer). The Ac and Su combination binds with 14/28 functional groups (Cytosolic Protein, Enzyme, Eraser, Family A GPCR, Hydrolase, Ion Channel, Kinase, Lyase, Membrane Receptor, Oxidoreductase, Phosphatase, Protease, Secreted Protein and Transferase). In our opinion considering the potential synergistic effects associated with use of ASwt in combinations, this should be avoided as it is likely to potentiate adverse effects.

A recent prospective cohort study revealed a positive association between ASwt consumption and atrial fibrillation (AF) rates,^27^ our study revealed potential mechanisms that may explain this association. We found that the targets of the ASwt we studied included several important proteins that have been found to be implicated in AF; KCNA5, KCNH2, TRPV4, BCL2, GSK3B. Although none of these targets were significant (C/A ratio ≤ 1.9), the chronic consumption of ASwt can lead to its accumulation in tissues niches, eventually raising to concentrations sufficient to activate these targets responsible for inducing AF. Also, the following high affinity targets of ASwt, CAPN1, LTA4H, MKNK2, ITGA2 and HDAC1 can indirectly regulate factors which can predispose to AF. These findings merit further studies, particularly ones that involve taking a chrono-pharmacological approach,^28^ to assess rates of AF events in relation to chronic ASwt consumption. We have previously examined chrono-pharmacology of other chronically used therapeutic and have demonstrated the association of periodic tissue accumulation and clinical presentation of adverse events.^38^ Such a chrono-pharmacological profile, merit following a dosing approach that allows for a washout phase to clear the active drug from the system to prevent adverse events occurring, consequence to the drug accumulating and building in tissue specific niche. Hence, based on this prior chrono-pharmacology knowledge we propose all chronic users of ASwt to allow for a few weeks (ideally 1-2 weeks) of washout phase every 6 months or alternatively to try a rotational use approach between Sa, Ac and St, with each ASwt being used for a few weeks sequentially.

The link between ASwt and cancer risk remains a subject of ongoing investigations. While some major regulatory bodies have deemed no convincing evidence for a direct cause-and-effect relationship, some studies suggest a possibility of associations between ASwt consumption and increased risk of developing cancer although without much insights into the mechanisms responsible.^13-15^ Our study addresses this gap in the literature by potentially identifying several ASwt targets, such as MCL1, ROCK2, BCL2L1, BCL2, MDM2, TP53, CDK proteins, HDAC1, ITGA2 and caspases which are widely reported to be associated with cancer development and/or progression.^39^ Incidentally high affinity targets of ASwt were highly expressed in endocrine systems, which again may highlight the increased risk of developing cancer. These ASwt targets are widely reported to influence a variety of oncogenes, tumour suppressor genes, extracellular matrix, apoptosis regulation proteins, and cell cycle regulating proteins.^39^ ASwt consumption has been specifically linked to increased risk of developing pancreatic cancer,^40^ whilst other studies have found pancreatic adenocarcinoma development to involve ROCK2 pathways,^41^ which we found to be a significant target of As, possibly underlining a mechanism through which As can lead to the development of pancreatic cancer. However, the potential interaction of ASwt, particularly As, with cellular targets identified in this study warrants further exploration to understand if these interactions could play a role in cancer development or progression. Long-term, well-designed epidemiological studies are crucial to definitively assess the potential association between chronic ASwt consumption and cancer incidence. In the meantime, adhering to the revised intake levels suggested in this study will be prudent.

A preclinical study has shown negative effects of ASwt on sperm quality. Studies on mice exposed to high doses of As observed reduced sperm parameters like motility, viability, and normal morphology. Additionally, these studies reported DNA fragmentation and decreased sex hormone binding globulin (SHBG) and testosterone levels.^29,30^ It’s important to note that these were animal studies with high doses, and it’s unclear if similar effects translate to humans at recommended intake levels. However, the findings raise concerns and warrant further investigation. While some studies have suggested a potential link between ASwt consumption and reduced fertility, particularly in women undergoing IVF (In Vitro Fertilization).^42,43^ Studies have shown that high intake of regular or diet soft drinks, containing ASwt, may be associated with decreased egg quality, embryo quality, and reduced implantation and pregnancy rates. The potential antifertility mechanisms could involve altered gut microbiome, disruption of hormonal pathways or directly targeting reproductive organs, all of which are crucial for sperm production and optimal functioning of gonads. In addition, this study highlights the potential role of SHBG in infertility associated with chronic consumption of ASwt, as SHBG was identified as a high affinity target of Su and both testis and ovary were observed to be pharmacodynamically significant tissue as these organs highly expressed significant targets ≥ 10 times of ASwt.

The potential link between ASwt and cardiovascular disease (CVD) is a topic of growing interest, with our findings adding a layer of complexity to this topic. While regulatory bodies generally consider ASwt safe at recommended intake levels, some observational studies suggest an association between high ASwt consumption and an increased risk of CVD.^5-8^ This study did reveal potential interactions between ASwt, particularly As and Su, with cellular targets (CAPN1, LTA4H, MKNK2, ITGA2, and HDAC1) involved in various physiological processes. Notably, some of these targets are linked to functions relevant to CVD development. For instance, the high C/A ratios observed for As with targets like ACE (Angiotensin Converting Enzyme) suggest a potential for influencing blood pressure regulation.^39^ Additionally, interactions with targets related to inflammation and cell death could also be relevant to CVD pathogenesis. The associations between ASwt and cardiovascular diseases may also be mechanistically explained by interactions with several targets, particularly: ACE, REN, AGTR1, HMGCR and NPC1L1. Influence of ASwt consumption on Hypertension^6,9,38^ can be explained by ASwt interactions with ACE, REN, AGTR1. Increased cholesterol uptake is a predisposing factor for many cardiovascular diseases, and this may be accounted for by interactions with HMGCR, and NPC1L1.^39^ Increased cholesterol uptake is associated with coronary artery disease, increased myocardial infarction risk, stroke, and atherosclerosis. The prevalence of these diseases has been correlated with consumption of ASwt.^44-46^ Future research should focus on randomised clinical trials with long-term follow-up to definitively determine if chronic ASwt consumption at recommended doses causally increases the risk of cardiovascular disease.

This study revealed potential interactions between ASwt, particularly As and Su, with various cellular targets at achievable doses. These interactions raise concerns about potential adverse health effects, especially in the gastrointestinal tract and closely associated organs, where some targets linked to inflammation (LTA4H)^47^ and cell death (CAPN1)^48^ were highly expressed. Furthermore, the high C/A ratios observed for some As and Su targets and their organ specific expression patterns suggest a possible increased risk of functional modulation in not only gastrointestinal tract but also endocrine, respiratory, renal, reproductive, central nervous, and musculoskeletal systems. We also observed colon to be a pharmacodynamically significant tissue impacted by ASwt, which may possibly explain observations regarding ASwt consumption and impacts on the gut micriobiota.^49^ ASwt interactions with the kidneys, which we also discovered to be a pharmacodynamically significant tissue, may explain associations with nephrotoxicity^50^ and chronic kidney disease.^16^

Despite some interesting insights into the pharmacodynamic effects of ASwt highlighted in this study, it does have some limitations. This study exclusively relied on in silico analysis, and hence in vivo trials are essential to validate these findings. Additionally, the long-term consequences of ASwt exposure require dedicated chrono-pharmacology focused research to establish a definitive link between consumption and potential health risks.

In conclusion, ASwt are widely used as sugar substitutes, but their impact on health remains a topic of concern. While considered generally safe at recommended doses by regulatory bodies, our findings suggest a need to exercise caution. Our study highlights the potential for ASwt to interact with various biological targets and induce adverse effects, particularly As and Su. The high C/A ratios of some As and Su targets and the tissue-specific expression patterns suggest potential safety concerns that require further investigation under long-term randomised settings.

## Acknowledgements

Research support from University College Dublin-Seed funding/Output Based Research Support Scheme (R19862, 2019), Royal Society-UK (IES\R2\181067, 2018) and Stemcology (STGY2917, 2022) is acknowledged.

## References

1. Chattopadhyay S, Raychaudhuri U, Chakraborty R. Artificial sweeteners–a review. Journal of food science and technology. 2014;51:611–621.

2. Sharma A, Amarnath S, Thulasimani M, Ramaswamy S. Artificial sweeteners as a sugar substitute: Are they really safe? Indian journal of pharmacology. 2016;48:237–240.

3. WHO R-LM, Montez J. Health Effects of the use of non-Sugar Sweeteners: a Systematic Review and Meta-Analysis. World Health Organization https://www.who.int/publications-detail-redirect/9789240046429. (Accessed on 15th March 2024) 2022.

4. Voiculescu DI, Ostafe V, Isvoran A. Computational assessment of the pharmacokinetics and toxicity of the intensive sweeteners. Farmacia. 2021;69:1032–1041.

5. Gomez-Delgado F, Torres-Peña JD, Gutierrez-Lara G, Romero-Cabrera JL, Perez-Martinez P. Artificial sweeteners and cardiovascular risk. Current Opinion in Cardiology. 2023;38:344–351.

6. Li B, Yan N, Jiang H, Cui M, Wu M, Wang L, Mi B, Li Z, Shi J, Fan Y. Consumption of sugar sweetened beverages, artificially sweetened beverages and fruit juices and risk of type 2 diabetes, hypertension, cardiovascular disease, and mortality: A meta-analysis. Frontiers in Nutrition. 2023;10:1019534.

7. Ebbeling CB, Feldman HA, Steltz SK, Quinn NL, Robinson LM, Ludwig DS. Effects of sugar-Sweetened, artificially Sweetened, and Unsweetened beverages on cardiometabolic risk factors, body composition, and sweet taste preference: a randomized controlled trial. Journal of the American Heart Association. 2020;9:e015668.

8. Diaz C, Rezende LF, Sabag A, Lee DH, Ferrari G, Giovannucci EL, Rey-Lopez JP. Artificially sweetened beverages and health outcomes: an umbrella review. Advances in Nutrition. 2023.

9. Qin P, Li Q, Zhao Y, Chen Q, Sun X, Liu Y, Li H, Wang T, Chen X, Zhou Q. Sugar and artificially sweetened beverages and risk of obesity, type 2 diabetes mellitus, hypertension, and all-cause mortality: a dose–response meta-analysis of prospective cohort studies. In: Eur J Epidemiol. 2020 Jul;35(7):655–671.

10. Alsunni AA. Effects of artificial sweetener consumption on glucose homeostasis and its association with type 2 diabetes and obesity. International journal of general medicine. 2020:775–785.

11. Imamura F, O’Connor L, Ye Z, Mursu J, Hayashino Y, Bhupathiraju SN, Forouhi NG. Consumption of sugar sweetened beverages, artificially sweetened beverages, and fruit juice and incidence of type 2 diabetes: systematic review, meta-analysis, and estimation of population attributable fraction. BMJ. 2015;351.

12. Malik VS, Li Y, Pan A, De Koning L, Schernhammer E, Willett WC, Hu FB. Long-term consumption of sugar-sweetened and artificially sweetened beverages and risk of mortality in US adults. Circulation. 2019;139:2113–2125.

13. Debras C, Chazelas E, Srour B, Druesne-Pecollo N, Esseddik Y, de Edelenyi FS, Agaësse C, De Sa A, Lutchia R, Gigandet S. Artificial sweeteners and cancer risk: Results from the NutriNet-Santé population-based cohort study. PLoS medicine. 2022;19:e1003950.

14. Mishra A, Ahmed K, Froghi S, Dasgupta P. Systematic review of the relationship between artificial sweetener consumption and cancer in humans: analysis of 599,741 participants. International journal of clinical practice. 2015;69:1418–1426.

15. Schernhammer ES, Bertrand KA, Birmann BM, Sampson L, Willett WC, Feskanich D. Consumption of artificial sweetener–and sugar-containing soda and risk of lymphoma and leukemia in men and women. The American journal of clinical nutrition. 2012;96:1419–1428.

16. Lo W-C, Ou S-H, Chou C-L, Chen J-S, Wu M-Y, Wu M-S. Sugar-and artificially-sweetened beverages and the risks of chronic kidney disease: a systematic review and dose–response meta-analysis. Journal of nephrology. 2021:1–14.

17. Gil TE, Gil JL, Laverde J. Artificially sweetened beverages beyond the metabolic risks: a systematic review of the literature. Cureus. 2023;15.

18. Christofides EA. POINT: artificial sweeteners and obesity—not the solution and potentially a problem. Endocrine Practice. 2021;27:1052–1055.

19. Yu B, Sun Y, Wang Y, Wang B, Tan X, Lu Y, Zhang K, Wang N. Associations of artificially sweetened beverages, sugar-sweetened beverages, and pure fruit/vegetable juice with visceral adipose tissue mass. Diabetes & Metabolic Syndrome: Clinical Research & Reviews. 2023;17:102871.

20. Sun Y, Yu B, Wang Y, Wang B, Tan X, Lu Y, Zhang K, Wang N. Associations of sugar-sweetened beverages, artificially sweetened beverages, and pure fruit juice with nonalcoholic fatty liver disease: cross-sectional and longitudinal study. Endocrine Practice. 2023.

21. Sadagopan A, Mahmoud A, Begg M, Tarhuni M, Fotso M, Gonzalez NA, Sanivarapu RR, Osman U, Kumar AL, Mohammed L. Understanding the role of the gut microbiome in diabetes and therapeutics targeting leaky gut: a systematic review. Cureus. 2023;15.

22. Pang MD, Goossens GH, Blaak EE. The impact of artificial sweeteners on body weight control and glucose homeostasis. Frontiers in nutrition. 2021;7:333.

23. Li D, O’Brien JW, Tscharke BJ, Choi PM, Ahmed F, Thompson J, Mueller JF, Sun H, Thomas KV. Trends in artificial sweetener consumption: A 7-year wastewater-based epidemiology study in Queensland, Australia. Science of the Total Environment. 2021;754:142438.

24. Silva Monteiro L, Kulik Hassan B, Melo Rodrigues PR, Massae Yokoo E, Sichieri R, Alves Pereira R. Use of table sugar and artificial sweeteners in Brazil: National Dietary Survey 2008–2009. Nutrients. 2018;10:295.

25. Huvaere K, Vandevijvere S, Hasni M, Vinkx C, Van Loco J. Dietary intake of artificial sweeteners by the Belgian population. Food Additives & Contaminants: Part A. 2012;29:54–65.

26. Daher M, Fahd C, Nour AA, Sacre Y. Trends and amounts of consumption of low-calorie sweeteners: A cross-sectional study. Clinical Nutrition ESPEN. 2022;48:427–433.

27. Basílio M, Silva LJ, Pereira AM, Pena A, Lino CM. Artificial sweeteners in non-alcoholic beverages: Occurrence and exposure estimation of the Portuguese population. Food Additives & Contaminants: Part A. 2020;37:2040–2050.

28. Heberle H, Meirelles GV, da Silva FR, Telles GP, Minghim R. InteractiVenn: a web-based tool for the analysis of sets through Venn diagrams. BMC bioinformatics. 2015;16:1–7.

29. Kumar AH. Network pharmacology of xylazine to understand its health consequences and develop mechanistic based remediations. bioRxiv. 2024:2024.2002. 2008.579475.

30. Kumar AH. Network Proteins of Human Sortilin1, Its Expression and Targetability Using Lycopene. Life. 2024;14:137.

31. Singh NK, Dhanasekaran M, Kumar AHS. Pharmacology of Berberine and Its Metabolites, Is It the Natures Ozempic or Imatinib? Arch Pharmacol Ther. 2023;5(1):67–81.

32. Friend J, Kumar AHS. A Network Pharmacology Approach to Assess the Comparative Pharmacodynamics of Pharmaceutical Excipient Trehalose in Human, Mouse, and Rat. Nat Cell Sci 2023;1(2):33–43.

33. Khosravi, Z., Kaliaperumal, C., and Kumar, A. H. S., (2023). Analysing the Role of SERPINE1 Network in the Pathogenesis of Human Glioblastoma. J Cancer Res. 1(1), 1–6.

34. Larsen JC. Artificial sweeteners: A brief review of their safety issues. Nutrafoods. 2012;11:3–9.

35. Rozman KK, Doull J. Paracelsus, Haber and Arndt. Toxicology. 2001;160:191–196.

36. Kim S, Chen J, Cheng T, Gindulyte A, He J, He S, Li Q, Shoemaker BA, Thiessen PA, Yu B. PubChem 2023 update. Nucleic acids research. 2023;51:D1373–D1380.

37. Blakemore E. Addicted to diet soda? Here’s the history of its low-calorie secret weapon. National Geographic. https://www.nationalgeographic.com/premium/article/aspartame-artificial-sweetener-health-history. (accessed on 17 March 2024). 2023.

38. Kumar AH, George M. Evaluating the Chrono-pharmacology of Icosapent Ethyl to Assess its Therapeutic Efficacy in Reducing Hypertriglyceridemia Associated Cardiovascular Events. Biology, Engineering, Medicine and Science Reports. 2019;5:26–28.

39. Hoda SA, Cheng E. Robbins basic pathology. In: Oxford University Press US; 2017.

40. McCullough ML, Hodge RA, Campbell PT, Guinter MA, Patel AV. Sugar-and artificially-sweetened beverages and cancer mortality in a large US prospective cohort. Cancer Epidemiology, Biomarkers & Prevention. 2022;31:1907–1918.

41. Liu J, Zhu Y, Ge C. LncRNA ZFAS1 promotes pancreatic adenocarcinoma metastasis via the RHOA/ROCK2 pathway by sponging miR-3924. Cancer cell international. 2020;20:1–15.

42. Halpern G, Braga D, Setti AS, Figueira R, Iaconelli A, Borges E. Artificial sweeteners-do they bear an infertility risk? Fertility and Sterility. 2016;106:e263.

43. Setti AS, Braga DPdAF, Halpern G, Rita de Cássia SF, Iaconelli Jr A, Borges Jr E. Is there an association between artificial sweetener consumption and assisted reproduction outcomes? Reproductive biomedicine online. 2018;36:145–153.

44. Zhang Z, Zhang K, Sun Y, Yu B, Tan X, Lu Y, Wang Y, Xia F, Wang N. Sweetened beverages and incident heart failure. European Journal of Preventive Cardiology. 2023;30:1361–1370.

45. Koeth RA, Smith JD, Chung M. Artificial Sweeteners: A New Dietary Environmental Risk Factor for Atrial Fibrillation? In: Am Heart Assoc; 2024:e012761.

46. Pase MP, Himali JJ, Beiser AS, Aparicio HJ, Satizabal CL, Vasan RS, Seshadri S, Jacques PF. Sugar-and artificially sweetened beverages and the risks of incident stroke and dementia: a prospective cohort study. Stroke. 2017;48:1139–1146.

47. DuBois RN. Leukotriene A4 signaling, inflammation, and cancer. In: Oxford University Press; 2003:1028–1029.

48. Tan Y, Wu C, De Veyra T, Greer PA. Ubiquitous calpains promote both apoptosis and survival signals in response to different cell death stimuli. Journal of Biological Chemistry. 2006;281:17689–17698.

49. Conz A, Salmona M, Diomede L. Effect of non-nutritive sweeteners on the gut microbiota. Nutrients. 2023;15:1869.

50. Ardalan MR, Tabibi H, Attari VE, Mahdavi AM. Nephrotoxic effect of aspartame as an artificial sweetener: A brief review. Iranian journal of kidney diseases. 2017;11:339.

